# In search of a dynamical vocabulary: a pipeline to construct a basis of shared traits in large-scale motions of proteins

**DOI:** 10.1101/2022.06.21.497011

**Authors:** Thomas Tarenzi, Giovanni Mattiotti, Marta Rigoli, Raffaello Potestio

**Affiliations:** Physics Department, University of Trento, via Sommarive, 14 I-38123 Trento, Italy; INFN-TIFPA, Trento Institute for Fundamental Physics and Applications, I-38123 Trento, Italy; Centre for Integrative Biology (CIBIO), University of Trento, Via Sommarive 9, 38123, Trento, Italy

## Abstract

The paradigmatic sequence-structure-dynamics-function relation in proteins is nowadays well established in the scientific community; in particular, large effort has been spent to probe the first connection, indeed providing convincing evidence of its strength and rationalising it in a quantitative and general framework. In contrast, however, the role of dynamics as a link between structure and function has eluded a similarly clear-cut verification and description. In this work, we propose a pipeline aimed at building a basis for the quantitative characterisation of the large-scale dynamics of a set of proteins, starting from the sole knowledge of their native structures. The method hinges on a dynamics-based clusterization, which allows a straightforward comparison with structural and functional protein classifications. The resulting basis set, obtained through the application to a group of related proteins, is shown to reproduce the salient large-scale dynamical features of the dataset. Most interestingly, the basis set is shown to encode the fluctuation patterns of homologous proteins not belonging to the initial dataset, thus highlighting the general applicability of the pipeline used to build it.

## I. INTRODUCTION

Internal large-scale motions of proteins are intimately linked to protein function [1]. These motions, happening on timescales of the order of *μ*s-ms [2], typically involve the collective fluctuations of secondary structure elements, leading to a variety of potential conformational states that might promote the exposure of specific binding sites [3, 4], or might facilitate the induced fit of the protein upon interaction with partner molecules [5, 6]. In addition, it has been shown not only that large-scale dynamics is essential for a protein to carry out its biological role [7], but also that a remarkable correlation exists between a proteins function and its specific dynamical signature [8], thus strengthening the view of dynamics as a link between a protein’s structure and its specific function. This is particularly evident for the case of allosteric proteins, where the binding of a ligand conveys a signal that is propagated within the protein structure through a modulation of its internal dynamics, resulting in alternative conformational states and an altered protein function [9–11].

Several computational methods exist for the study of slow dynamics in proteins [12–14]; however, in order to develop a more general view of how dynamics bridges structure and function, it is necessary to build a dataset-wise approach for the comparison of such large-scale dynamics among proteins sharing different degrees of sequence and structural similarity. Attempts in this direction have been performed in several works [15–20]. Maguid et al. [21] based their analysis on a dataset of pairs of homologous proteins; comparison of vibrational backbone dynamics within each pair led to the remarkable observation of correlation between dynamics and evolutionary conservation. Analyses of the distance in dynamics have also been performed in the case of structurally and functionally diverse sets of proteins; in this regard, Hensen et al. [8] introduced the notion of “dynasome”, namely an ensemble of observables computed from molecular dynamics (MD) simulations of a structurally heterogeneous protein dataset. The method highlights a striking correlation between the dynasome descriptors (which include 34 observables for each protein, ranging from the first five eigenvalues of the covariance matrix of C_*α*_ fluctuations to the average ruggedness of the energy landscape) and the proteins functional classification. However, this approach relies on time-consuming MD simulations, which limits its applicability to large protein datasets. In addition, the large number and sophistication of the descriptors employed does not enable a straightforward recognition and visualization of the similarities in dynamics between proteins in term of conformational movements.

To overcome these limitations, in this work, we setup and validate a novel pipeline for the identification of a basis set of conformational motions in an enzymatic family, representing a common vocabulary of their large-scale dynamics. To this aim, we investigated internal protein dynamics in terms of fluctuations at the level of single residues. Our approach does not require the acquisition of expensive MD simulations, since it is based on the topology of native contacts derived from a protein’s experimental structure; specifically, we made use of the normal mode analysis (NMA) [22], which represents, together with the principal component analysis (PCA) [23], one of the main protocols employed to identify the most relevant patterns in the large-scale dynamics of proteins. While PCA requires a large set of configurations (for example from MD trajectories) to build the covariance matrix, NMA can be performed with the sole knowledge of an equilibrium configuration of the system. For this reason, NMA is often used in combination with simplified quadratic models, such as the linearized versions of elastic network models (ENMs) [24]. Another degree of simplification can also be introduced by building coarse-grained (CG) models of the protein, where the atomistic degrees of freedom are replaced by a smaller number of physically relevant representative beads. In spite of this simplicity, the collective, large-scale dynamical features obtained by NMA of ENMs of proteins showed to be successful to predict experimental B-factors [25] and also conformational changes [26, 27].

In our approach, normal modes are computed from the *β*-Gaussian elastic network model of a set of chymotrypsin-related proteases, for which in-depth analyses of evolutionary relationships and structural similarities are available in the literature [28–31]. In the *β*-Gaussian model, each residue is described in a simplified representation as two beads [32]: one corresponds to the C_*α*_ atom and represents the mainchain, while the second, describing the sidechain, is positioned according to the degrees of freedom of the first bead. An effective quadratic potential energy is used to model the bead fluctuations from the native conformation. We made use of this information to perform a dynamics-based alignment between all pairs of proteins from the dataset; the results from the alignment were used to construct a distance matrix in the space of protein dynamics and to cluster together proteins with similar large-scale motions, thus adding an additional layer of information to clustering procedures based on sequence identity [33, 34] or structural similarity [35–37].

Moreover, we developed a way to represent each protein’s large-scale normal mode as a vector field on the 3D space. Thanks to this representation, we were able to build a high-dimensional basis set of large-scale protein modes. The basis set is validated by comparison with results from MD simulations, with the perspective of applying this methodology to a dataset comprehensive of a large number of protein classes, differing in structure and function. In this way, common fluctuations between distant proteins can be correlated to the presence of local structural elements, with implications in protein engineering for the design of scaffolds that are able to perform controlled conformational changes in functional enzymes [38, 39]. In addition, the large-scale dynamics might serve as a guide in the identification of those patterns where the preservation of a high resolution is of paramount importance in the construction of simplified, multiscale models [40–44] that retain the original dynamics. In particular, by describing at an atomistic level the structural elements identified as important for the desired conformational movements and simultaneously coarse-graining the remainder of the protein, it might be possible to obtain a simplified and computationally inexpensive protein model that shows the conformational dynamics of the high-resolution one.

The article is structured as follows: in Section II, the workflow developed for the construction of the basis set of conformational moves is described; in Section III, technical details on the methods employed are given; in Section IV, the results are presented and discussed; eventually, the conclusions and the future perspectives are discussed in Section V.

## II. OVERVIEW OF THE WORKFLOW

In our approach, the identification of a common set of conformational motions among different proteins is based on the analysis of their dynamics in a CG representation; from here, a representative set of normal modes is identified through a dynamics-based clustering of the proteins comprising the initial dataset. The selected, representative modes are then orthonormalized and ordered, so as to obtain the final basis set. An overview of the workflow is given in Figure 1 and explained in detail in the following paragraphs.

**FIG. 1:**
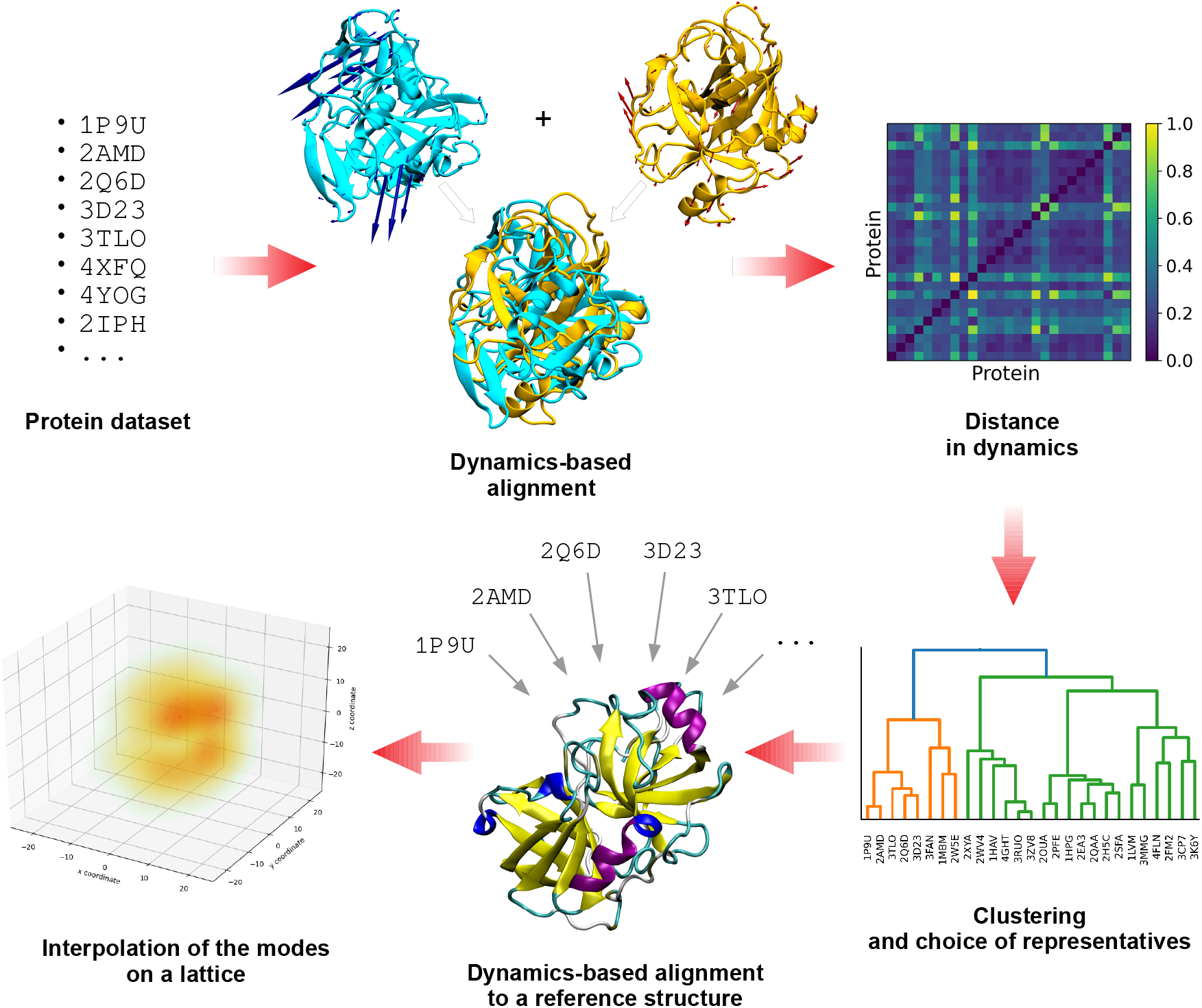
Schematic representation of the workflow proposed, from the choice of the protein dataset to the creation of the vector fields on a grid. Once orthonormalized and ordered, the latter are used to construct the final basis set.

The starting point is the identification of a set of proteins. The choice of this dataset is arbitrary and independent on the pipeline; however, the number of proteins that the dataset contains is supposed to be large enough so as to be representative of the families or superfamilies that are included, meaning that the more distant are the members in terms of homology, the larger should be the dataset. This is necessary to ensure sufficient generality of the resulting basis set of conformational motions.

The selected set of structures is used to run pairwise dynamics-based protein alignments with the ALADYN software developed by some of us [45]. ALADYN takes two input structures and performs the maximization of a score function that takes into account the spatial superposition of protein regions that have similar motion. The dynamical information is encoded in the low-energy (large-amplitude) eigenvectors obtained from the diagonalization of the interaction matrix *M*_*ij*_ of the Hamiltonian function of the *β*-Gaussian Network Model:

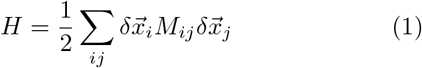

where 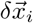 is the displacement vector of the i-th bead with respect to the equilibrium configuration. Once the eigenvectors have been obtained, the extent and consistency of the alignment are quantified through the root-mean-square inner product (RMSIP) between the spaces given by the first 10 modes of each aligned protein. If we call *N*_*i*_ and *N*_*j*_ the total number of residues in the chains of the two aligned proteins, the RMSIP calculation is limited to a subset *q < N*_*i*_, *N*_*j*_ of marked C_*α*_. These subset of amino acids are chosen by firstly grouping the amino acids in groups of 10 subsequent ones; then maximizing a single scoring parameter via the standard Metropolis criterion over the space of possible pairs of groups among the two proteins’ sequences, as exhaustively explained in [45]. Specifically, the RMSIP is defined as:

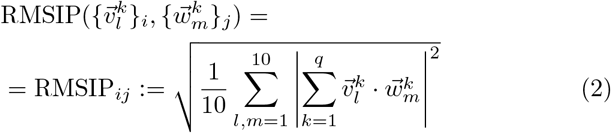

The RMSIP ∈ [0, 1] takes on the value 1 in case of perfect correspondence of the spaces, and 0 in case of their complete orthogonality. The quantity (1.0 - RMSIP), which still takes values in the interval [0, 1], is therefore suitable to define a distance in dynamics between two proteins after alignment. Statistical significance of the alignment, quantified by means of a *z*-score, is taken into account by weighting the RMSIP by the hyperbolic tangent of the module of the *z*-score, so as to give more importance to the most reliable results. The distance in dynamics between two aligned proteins *i* and *j* is therefore defined as:

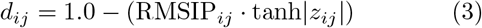

After all the pairwise alignments between the elements of the dataset are performed, a distance matrix that expresses differences in the large-scale dynamics is obtained; then the dataset undergoes hierarchical clustering [46] based on this distance matrix, in order to identify groups of dynamics-related proteins. The optimal number of clusters is identified from the interplay between *resolution* and *relevance* [47–51]. These two quantities, which are defined in more detail in the Methods section, are entropies that are related to each other and depend on the clusterization procedure adopted. We exploited them to select the number of clusters to retain, by considering the smallest number of clusters (hence the lowest resolution) that gives the highest relevance (Figure 2).

**FIG. 2:**
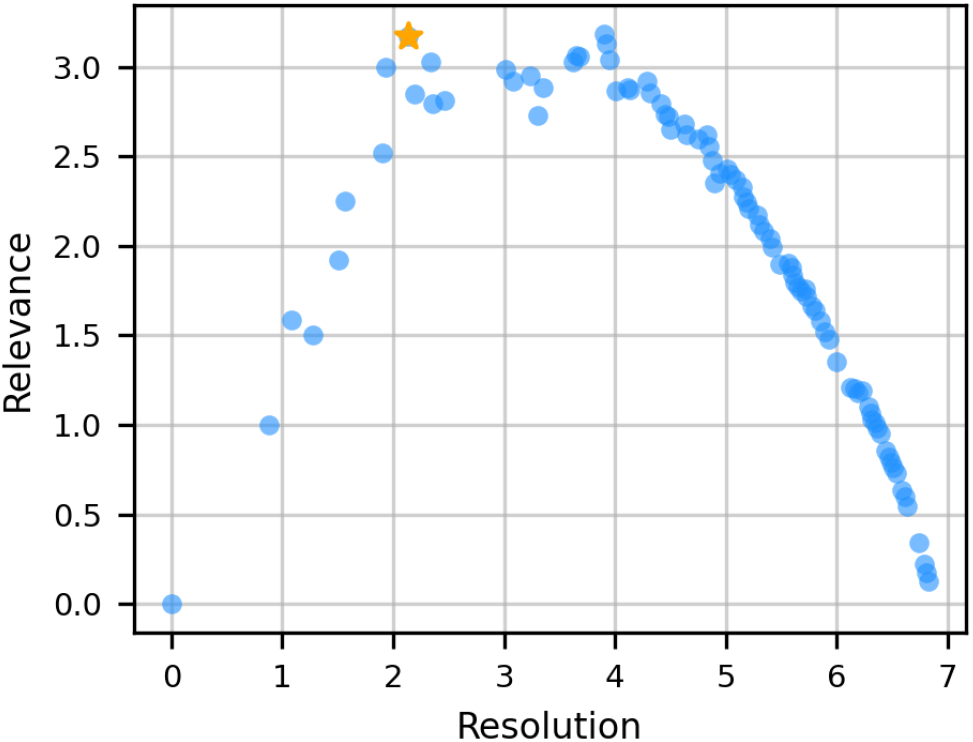
Relevance-resolution curve used to determine the optimal number of clusters. Each point corresponds to a different number of clusters. The optimal subdivision, indicated with an orange star, corresponds to 9 clusters.

Once the optimal number of clusters is derived, protein representatives of each cluster are identified as the cluster centroids, namely the proteins with the shortest distance to every other protein of the cluster itself. In addition, a representative for the whole dataset is selected as the protein with the most characteristic dynamics, expressed in terms of the lowest distance with respect to all the other dataset members. The other protein structures are then dynamically aligned to this one with ALADYN, so as to have a consistent orientation in space.

From an ENM representation of each of these newly oriented structures, normal modes are computed. In order to facilitate the comparison between modes belonging to proteins with a different sequence length, the first 5 reoriented normal modes of the cluster representatives are placed on a cubic lattice, and interpolated on the grid points so as to obtain a smooth vector field (Figure 1). In this way, we move from comparing the 3*N* -dimensional modes of different proteins (where *N* is the number of residues, different for each protein), to comparing vector fields defined on identical 3D lattices having the same dimension. More details on the lattice construction and interpolation are given in Section III. Proteins belonging to the dataset employed in this work, despite displaying a range of sequence length and radius of gyration, do not grandly differ in size; therefore, the modes interpolated on the lattice can be directly compared. However, it might be the case that the dataset includes proteins with very different size; this would require a rescaling of the protein coordinates before the interpolation on the lattice, so as to compare motions occupying similar volumes in space.

The interpolated modes are orthonormalized using the Gram-Schmidt algorithm [52]. The components of the basis are finally ordered according to decreasing entropy, considered as a measure of their degree of collectivity. The entropy *S* of a mode *k* is defined as:

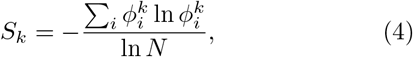

where *N* is the number of lattice sites and 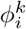 is the square modulus of the *k*-th mode on the lattice site *i. S*_*k*_ takes a maximum value of 1 if the mode is delocalized on all the lattice sites, and a minimum value of 0 if the mode is localized on a single site.

The final set of orthonormalized and ordered vector spaces represents the basis of protein dynamics. In the next section, technical details of the methods employed are presented.

## III. MATERIALS AND METHODS

### A. Preprocessing of the dataset

A dataset of 116 chymotrypsin-related proteases, for which structural experimental information is available, was selected. This dataset is based on the one used in ref. [31], from which proteins with sequence identity *>* 70% were removed. The dataset comprises serine proteases from bacteria, eukaryotes, archaea, and viruses, in addition to chymotrypsin-related cysteine proteases from positive-strand RNA viruses. The full list of proteins’ PDB IDs is given in Table S1. The structures were downloaded from the Protein Data Bank, and the co-ordinate files were cleaned-up from heteroatoms, from copies of the protein in the crystallographic cell, and from residue-configurations with low occupancy. The position of missing atoms was rebuilt and the protein conformations were optimized using the software FoldX [53]. Non-terminal missing residues were modelled with MODELLER [54, 55]. An analysis of the first 3 normal modes for each protein was run using an elastic network model with a cutoff of 10 Å, in order to identify the problematic cases in which the flexible protein termini impaired the analysis of the motion of the protein core. Such analysis was conducted by visual inspection of the modes on the protein structures. In those cases, flexible tails were not considered in the following analyses, which thus focused on globular structures. Moreover, in the case of multi-domain structures, only the domain known to have protease activity was retained.

### B. Dynamics-based alignment and clustering

The dynamics-based alignment of all the pairs of protein structures was performed with the ALADYN software [45], using as input the cleaned coordinates files. From the resulting alignment scores, clustering of the structures was performed with the Python library SciPy, using the ward linkage method.

The calculation of relevance and resolutions, used to identify the optimal number of clusters, was performed with an in-house script. Specifically, given a labeling *ŝ* := (*s*_1_, …, *s*_*η*_) (e.g. a clustering) to a sparse dataset made by *N* ≥ *η* data points (in our case the single proteins in the dataset), resolution is defined as an entropy *Ĥ*_*res*_ representing the relative amount of information loss in the process:

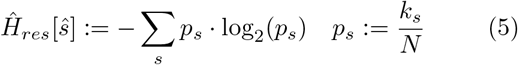

where *k*_*s*_ is the number of data points that fall into the same cluster *s*. It is proven [48] that *Ĥ*_*res*_ increases monotonically with the number of clusters, in accordance with the idea that the coarser is our clustering, the more information we loose. On the other hand, the relevance *Ĥ*_*rel*_ is defined as:

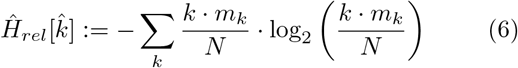

where *m*_*k*_ is the number of clusters containing the same amount *k* = 0, …, *N* of data points, for a given clustering process. By choosing the lowest resolution value corresponding to the largest relevance, we can rely on the most compact clusterization (thus increasing the statistics within each cluster) that preserves the highest empirical information content.

### C. Lattice interpolation and basis construction

Normal modes of each protein of the dataset have been computed with an in-house code. The first 5 reoriented normal modes of the cluster representatives were placed on a cubic lattice, with a lattice constant of 1 Å(for a total of 45 modes, namely vector fields). The vector on each protein C_*α*_ was translated on the nearest lattice grid point. The mode vectors were interpolated on the lattice in order to create a smooth vector field (Figure 1), using Gaussian functions with *σ*=0.8 Å and truncated at a distance of 2 Å. This distance is slightly smaller than the lowest spatial distance between two C_*α*_ atoms to make sure that the vector coming from the original protein mode is not spuriously modified during interpolation. The chosen value of *σ* ensures that in correspondence of the cutoff the mode field is close to zero. The resulting vector at each grid point *ijk* is the sum of the mode fields centered on the nearby C_*α*_ grid points, calculated at *ijk*, within the cutoff. Eventually, orthonormalization and ordering of the modes was performed with Python scripts.

### D. Molecular dynamics simulations

Molecular dynamics simulations have been performed on the representatives of each cluster, using the software Gromacs 2019 [56]. The proteins were described with the Amber99sb-ildn force field [57], and the TIP3P model [58] was used for water molecules. Sodium and chloride ions were added at a concentration of 0.15 M, and balanced so as to neutralize the charge in the simulation box. All systems were energy minimized for 100 steps by steepest descent. The solvent was then equilibrated for 500 ps with positional restraints on the protein heavy atoms, using a force constant of 1000 kJ·mol^−1^·nm^−2^. MD simulations were carried out in the NPT ensemble for 250 ns for each system. Protein and solvent were coupled separately to a 300 K heat bath with a coupling constant of 0.1 ps, using the velocity-rescaling thermostat [59]. The systems were isotropically pressure-coupled at 1 bar with a coupling constant of 2.0 ps, using the Parrinello-Rahman barostat [60]. Application of the LINCS [61] algorithm on hydrogen-containing bonds allowed for an integration time step of 2 fs. Short-range electrostatic and LennardJones interactions were calculated within a cut-off of 1.0 nm, and the neighbor list was updated every 10 steps. The particle mesh Ewald (PME) method was used for the long-range electrostatic interactions [62], with a grid spacing of 0.12 nm.

The calculation of the root-mean-square fluctuations from the trajectory coordinates was performed on the protein C_*α*_ atoms using the Gromacs tool *gmx rmsf*. The dynamic cross-correlation was computed with a Python script, using the library MDTraj [63]. Plots were produced with Python libraries, and protein images were rendered with VMD [64].

## IV. RESULTS AND DISCUSSION

### A. Overview of the protein dataset

Proteases are enzymes catalyzing the reaction of hydrolysis of peptide bonds. The independent evolutionary origin of these proteolytic enzymes, which led to a variety of chemical solutions for the same functional problem [65], is reflected in the large variety of sizes, shape and specificity of proteases [66]. In this work, however, we focus on a specific superfamily of proteases. The 116 members of the selected dataset are indeed chymotrypsin-related proteases, sharing a common structure with two *β*-barrel-like domains that accommodate the binding site (Figure 3). However, the length of the turns and loops connecting the sheets greatly varies; moreover, *α*-helices are present in some of the structures, as a result of adaptation to specific functions and recognition/binding of specific ligands. The result of this structural variability is a range of sequence lengths and protein sizes (Figure S1). Not only the number and type of secondary structures can vary, but also the size and structural completeness of the *β*-barrels themselves; for example, the 2A proteases of enteroviruses (family C3, subfamily C3B) have only four antiparallel *β*-strands in place of the N-terminal barrel [67]. In all the proteins included in the dataset, the proteolytic reaction is performed by a catalytic triad of residues, located between the *β*-barrels. The type of amino acid playing the role of nucleophile in the mechanism of catalysis determines the class of proteases: in the serine proteases, the catalytic triad contains His, Asp/Glu, and Ser residues [68]; in the cysteine proteases, the triad is composed of His, Asp/Glu, and Cys or of a dyad of His and Cys residues, as in the hepatitis A virus 3C protease and in the coronavirus 3C-like proteases [69]. The classification used in the remainder of the paper is based on MEROPS, a hierarchical classification scheme for proteases [70, 71]. In the MEROPS database, chymotrypsin-related proteases constitute the PA clan, which contains 9 families of cysteine proteases (representing proteases of positive-strand RNA viruses) and 14 families of serine proteases (representing proteolytic enzymes from eukaryotes, bacteria, some DNA viruses and eukaryotic positive-strand RNA viruses). Families are defined on the basis of sequence similarity and/or resemblance of the folds among their protein members. However, experimental structural information is available for a limited number of these families; therefore, not all of them are represented in the dataset employed in this work.

**FIG. 3:**
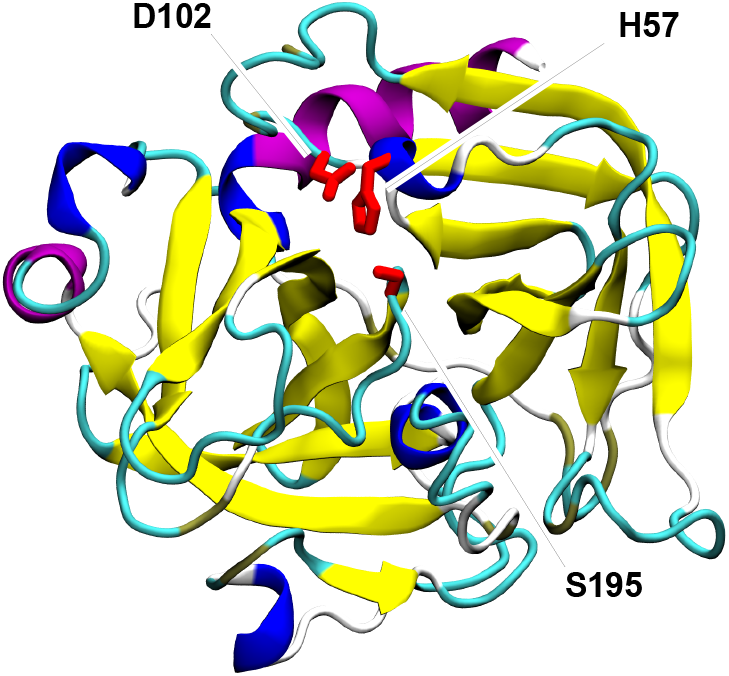
Cartoon representation of chymotrypsin from *Bos taurus* (PDB ID: 2CGA). Colors are used to differentiate the structural elements; in particular, the two *β*-barrels are distinguishable in yellow. The catalytic triad is represented in licorice and colored in red.

### B. Results of the dynamics-based alignment

We performed an alignment based on the dynamical information entailed into the first 10 lowest frequency modes obtained by the NMA on the *β*-Gaussian Network Model of each pair of proteins in the dataset. The alingment consists in the optimization of a score function that maximizes the RMSIP of the two sets of normal modes. The distance matrix obtained from the pairwise dynamics-based alignments of all proteins of this dataset is used as a measure of similarity in dynamics. This can be compared to the MEROPS classification by computing the average distance between protein pairs that fall into the same family. Following such a procedure, it is apparent that the average distance in dynamics is lower within each family, with respect to the total average (Figure 4). In other words, proteins are significantly closer in dynamics within the same family then they are to members of other families.

**FIG. 4:**
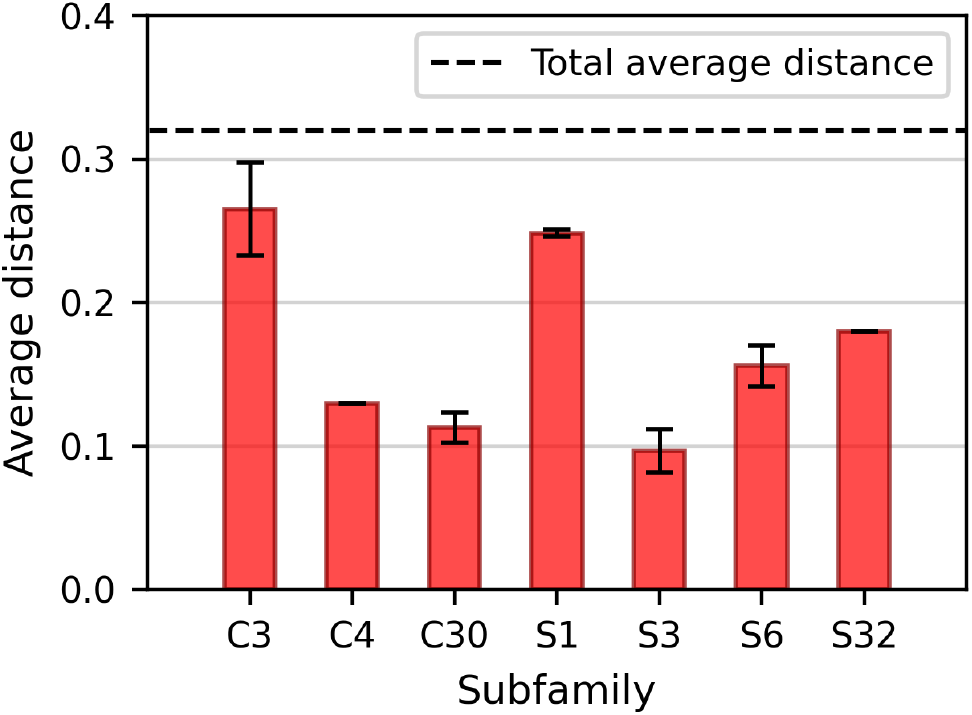
Average distances (in terms of dynamics) between proteins of the dataset belonging to the same family. Only those subfamilies including more than one representative member are displayed here.

The distance matrix is used as input for the division of the dataset into dynamically homogeneous protein clusters. The outcome of the hierarchical clustering is graphically expressed by the dendrogram in Figure S2. On the basis of the resolution-relevance plot, 9 clusters were identified (Figure 2); this corresponds to a threshold of ≈ 0.58 in the clustering dendrogram. The resulting clusters appear to be quite homogeneous in terms of protease classification (Figure S3). Importantly, the dynamics-based clustering automatically tends to group proteins belonging to the same subfamily. Figure 5.a shows that in most of the cases (17 of the 19 subfamilies represented in the dataset) all the members of each subfamily fall into the same cluster, thus suggesting that these proteins share a similar conformational dynamics and strengthening the idea of homogeneity in dynamics between homologous proteins [72, 73]. On the other hand, each cluster groups several subfamilies, and only 4 clusters out of 9 include proteins belonging to only one subfamily (Figure 5.b). Therefore, the clustering procedure proves able to effectively group different protein subfamilies that, despite the different evolutionary origin, share similar dynamics.

**FIG. 5:**
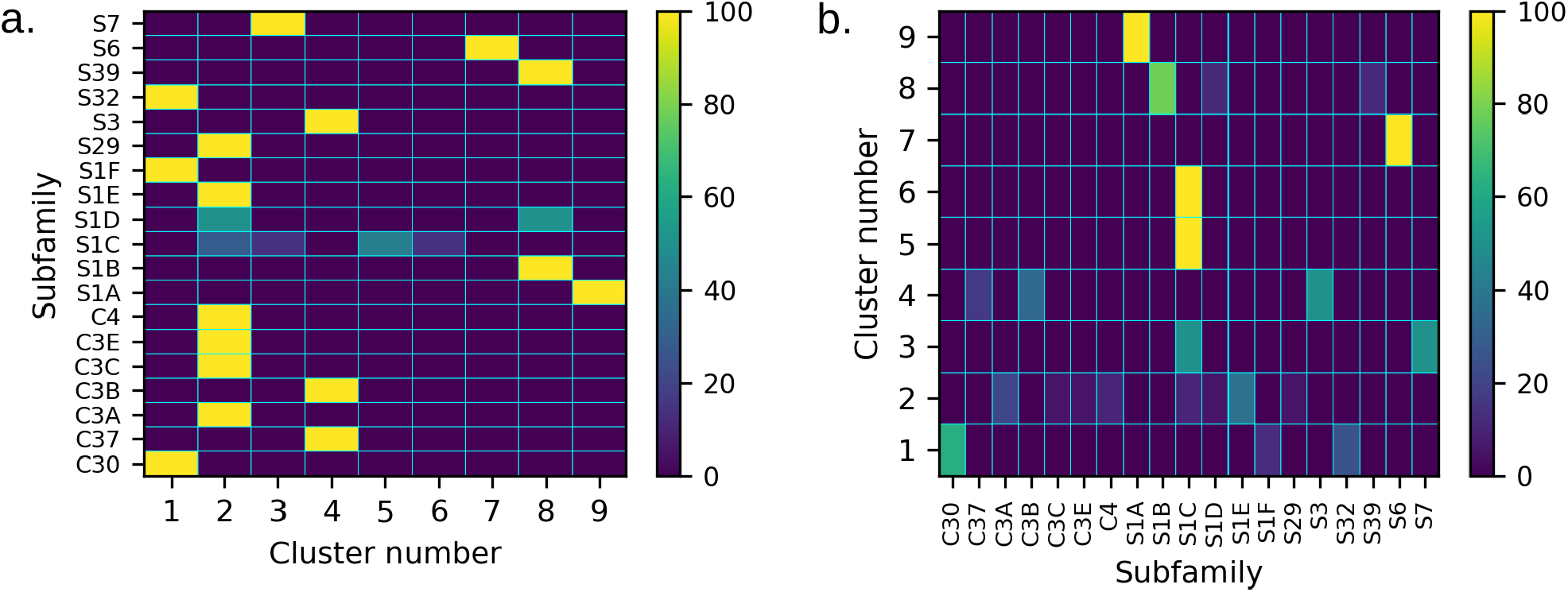
**a**. Distribution of the members of each subfamily among the different clusters, expressed as percentage with respect to the total number of members of the subfamily. In **b**. each row represents the content of each cluster classified on the basis of the function (in percentage with respect to the total population of the cluster).

### C. Comparison between the dynamics-based and the structure-based clustering

We compared the results from the dynamics-based clustering on the proteases of the PA clan with the structure-based distance tree calculated in the work of Mönttinen et al. [31]. There, the authors identified a common structural core of 72 residues for the set of PA clan proteases taken into account; according to the structural similarities of this common core, they built a distance tree between the members of the dataset. 5 different clusters were identified, contrary to the 9 cluster found in this work.

Despite the two different approaches, the results present several similarities, showing a close relation between structure and dynamics. The S1A subfamily, which includes both bacterial and eukaryotic proteases, forms a clearly distinct and compact cluster both in terms of structure and dynamics. On the other hand, the S1D subfamily, which includes bacterial proteases, is split in two different groups in terms of structure as well as dynamics: in both cases, the S1D *Achromobacter* protease I (1ARB) is close to the bacterial S1B proteases, while the S1D protease AL20 of *Nesterenkonia abyssinica* (3CP7) is close to the members of the bacterial S1E subfamily. This difference between members of the S1D subfamily has been explained on the basis of the different evolutionary history of the bacteria in which they are expressed [31].

Another common feature emerging from the two clustering approaches is the similarity between the S39 sub-family of positive-strand RNA viruses and the bacterial S1B proteases; interestingly, such degree of similarity is higher than between S39 and the other viral proteases, as already reported on the basis of structural comparisons [74]. Moreover, the bacterial S6 family forms an independent group in both clustering approaches. This peculiarity has been attributed to the presence of a long *β*-stalk structure at the C-terminus (Figure S4), which is absent in all the other proteases of the PA clan [31, 75]; the protease domain alone, instead, shares high structural similarity with that of the S1A subfamily. However, the *β*-stalk domain was cut before the dynamics-based alignment, meaning that our analysis of dynamics of the S6 protease domain alone is able to distinguish this subfamily from the other members of the PA clan.

Importantly, the two types of clustering present also some differences. In the case of the structure-based analysis, the cysteine proteases tend to be grouped together; however, in the dynamics-based alignment, the similarity is only at the level of one of the two large groups in which the dataset is divided, as evident from the dendrogram in Figure S3. Within this group, C families are mixed with S families, and appear to be more distributed among different clusters than in the distance tree built on the basis of the structural features. This is indicative of a clear differentiation of the C proteases in terms of dynamics, despite their structural similarity in the protein core. Another difference regards the heat-shock proteases S1C, which includes proteins from bacteria, chloroplasts, and mitochondria; even though structurally similar in the proteolitic core, members of this subfamily appear very scattered in the dynamics-based clustering. Specifically, the observed similarities in dynamics accentuates the structural relatedness already observed between some eukaryotic S1C proteases and different viral protease subfamilies, inasmuch that these similarities are stronger than the similarity within the S1C subfamily itself. This relatedness has been previously explained on the basis of exchanges of protease genes between eukaryotic viruses and their hosts [31].

In the structure-based distance tree, proteases from flavivirus (families S29 and S7) and from togavirus (family S3) are grouped together, even though the two viruses belong to different families; on the opposite, S29/S7 and S3 are placed in different clusters when their dynamics is included in the analysis. This distinction might arise from the difference in function: the S3 protein togavirin, in fact, does not only function as a viral protease, but plays also the structural role of capsid protein of the virus [76]. S29 and S7 proteases, on the other hand, possess only proteolitic function and do not work as structural components.

Overall, the inclusion of dynamics in the comparison of the proteases from the PA clan adds therefore an additional level of classification, which seems appropriate to bridge structural and functional similarities.

### D. Creation and validation of the basis set of the high-dimensional space of protein dynamics

The representative proteins of the 9 clusters are identified by the PDB codes: 3D23, 1HPG, 2YOL, 1VCP, 3QO6, 1L1J, 1WXR, 4JCN, 4I8H. Their structures are represented in Figure S5. Protein 1GDQ is chosen as the reference structure of the whole dataset, against which the other representatives are dynamically aligned prior to lattice interpolation of their normal modes (see Section III). In the latter, the oriented protein modes are placed and interpolated on a cubic lattice, orthonormalized, and finally ordered. The interpolation on the grid allows us to easily compare the dynamics of any pair of proteins, irrespectively on the number of residues. For instance, modes from proteins with a different number of C_*α*_ cannot be directly compared in terms of scalar products, while different vector fields on the grid have the same dimensionality.

We investigated the quality of the orthonormalized modes as a basis set for the dynamics of the whole dataset, by computing the overlap between the spaces given by the protein modes and by the basis. To this aim, the RMSIP was computed between the space spanned by the first 5 modes of each protein in the dataset (after their interpolation on the lattice) and the first *n* components of the basis. For each protein, the components of the basis are ordered so as to maximise the RMSIP with the protein modes. The resulting RMSIP for each protein is plotted in Figure 6.a as a function of the number *n* of basis vectors considered for the calculation of the RMSIP. From the distribution of the values attained when using the full basis set (45 vector fields), the RMSIP is greater than 0.5 for ≈ 94% of the proteins, showing in those cases a good agreement between the dynamics of the protein and the one expressed by the basis [77]. The agreement is excellent (RMSIP*>* 0.7) for ≈ 61% of the proteins; therefore, we can conclude that the identified basis is indeed able to describe with a good generality the large-scale conformational dynamics of the dataset. For each protein, we also computed the normalized RMSIP, by dividing each value of RMSIP with the value obtained with the use of the full basis set. The normalized RMSIP curves show that, for each dataset member, as few as 15 basis components are sufficient to reproduce the 80% of the dynamics that would be attained with the use of the full basis set (Figure 6.b); however, such components differ from protein to protein, meaning that there are no vector fields in the basis that can be considered more essential than others. This suggests that a further reduction in the dimension of the basis set would lead to a loss of generality in the description of the dynamics of this class of proteins.

**FIG. 6:**
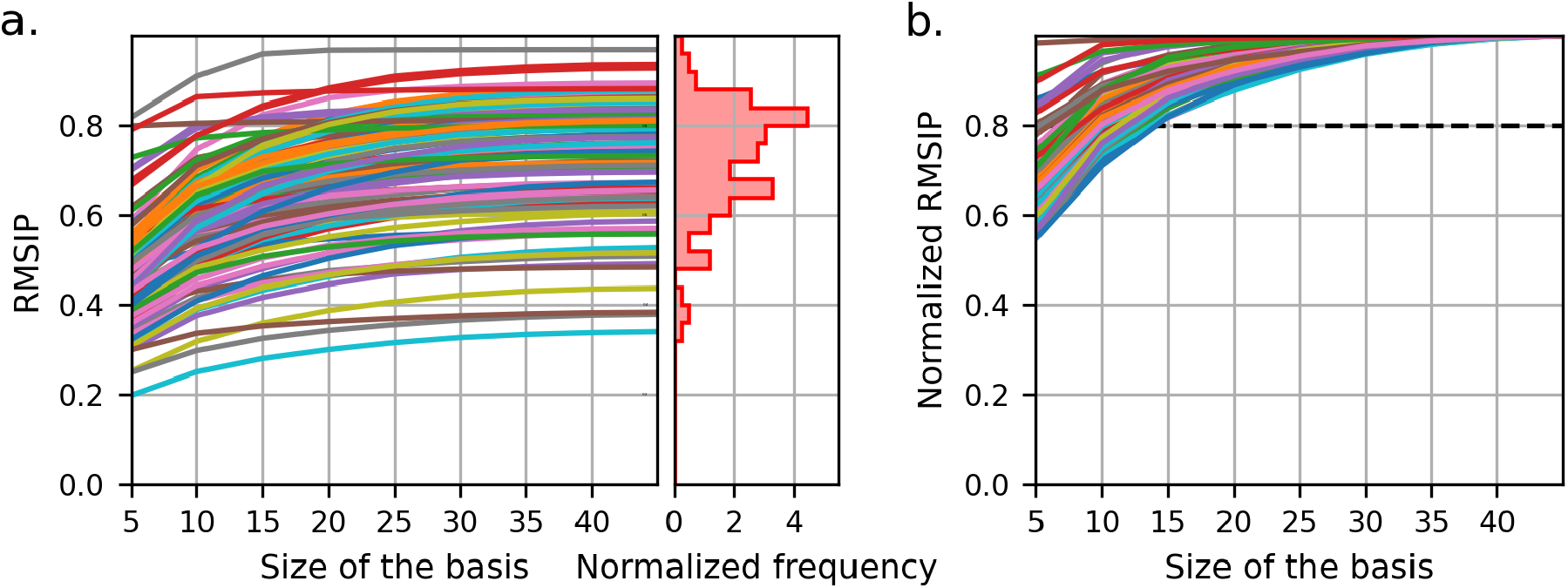
**a**. RMSIP between the subspaces spanned by the first 5 modes of each protein and the first *n* basis vectors, as a function of the basis size *n*. The histogram on the right represents the distribution of the RMSIP values attained when the full basis is used. **b**. RMSIP normalized with respect to the value attained from the use of the full basis.

### E. Comparison with MD simulations

In order to better assess the ability of the basis to reproduce the general dynamics of chymotrypsin-like proteases, we performed MD simulations of four proteins belonging to the same family, and compared the per-residue fluctuations emerging from the simulations with those obtained by filtering the trajectory along the vectors of the basis; a good agreement would be indicative of the ability of the basis to describe the large-scale dynamics of the protein. Two of the proteins used as test-case belong to the dataset; these are 1EKB [78] and 1NPM [79], eukaryotic proteases belonging to the S1A subfamily. The other two proteins, 4YOG [80] and 3W94 [81], are external to the dataset, and as such have not been used to define the basis. 4YOG is a C30 protease from the bat coronavirus HKU4, while 3W94 is an S1A enteropeptidase. These two proteins have been included here in order to test the generality of the identified basis for the description of the dynamics of the PA clan, independently on the specific members of the initial dataset.

For each of the four proteins we compared the root-mean-square fluctuations (RMSF) as computed from the simulation, and as computed from the same trajectory filtered along the modes given by the backmapping of the protein structure on the basis vectors. The comparison shows a good qualitative agreement (Figure 7 and Figure S6), in particular in correspondence of all the secondary structure elements. In the unstructured regions, the comparison is slightly less accurate; this is particularly true for long loops, which are more sensitive to the limitations of the ENM and of the NMA employed to define the modes of the basis, since both assume small-amplitude fluctuations from a well-defined reference structure. From the two sets of trajectories, namely the original MD simulations and the filtered ones, we also computed the dynamic cross-correlation matrices (Figure S7 and Figure S8), which give a measure of the degree of correlation between each pair of C_*α*_ atoms in terms of fluctuations from their average position. When comparing the original and filtered trajectories, the intensity of the resulting correlations are different, with higher correlations/anti-correlations emerging form the trajectory filtered on the basis; however, the patterns of correlation are strikingly similar between the two trajectories for all of the four proteins. Also in this case, therefore, the basis set appears to be able to describe the relevant large-scale dynamics of the considered protein systems.

**FIG. 7:**
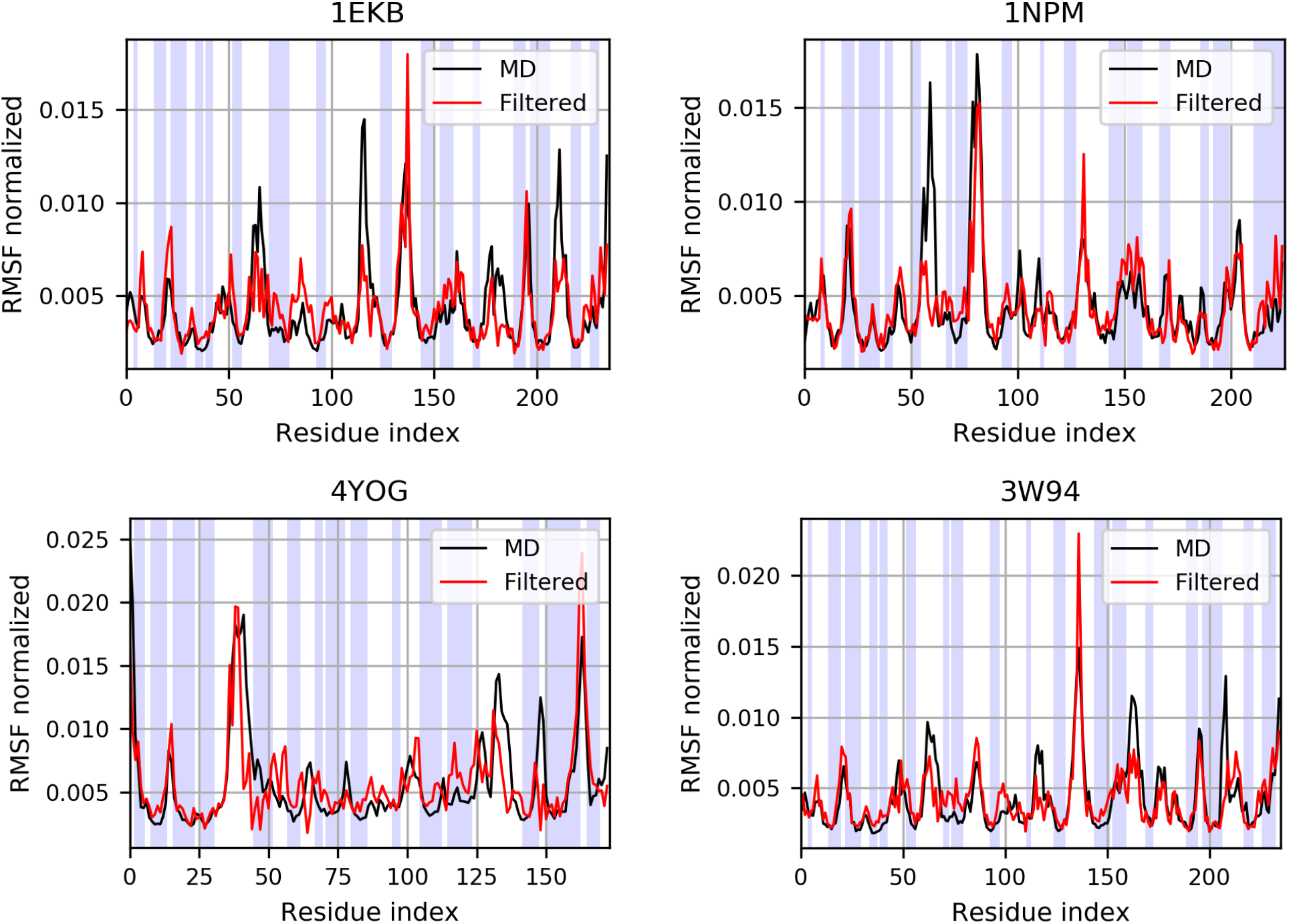
Root-mean-square fluctuations of the C_*α*_ atoms, normalized with respect to their sum for a qualitative comparison. The shaded areas correspond to structured regions of the protein, identified with the DSSP algorithm [82, 83].

## V. CONCLUSIONS

In this work, we proposed a workflow for the identification of common large-scale conformational motions in a set of proteins. Specifically, we performed a dynamics-based clusterization of 116 chymotrypsin-related proteases, belonging to the PA clan, and compared the resulting clusters to the MEROPS classification and to a more recent structure-based classification of the same dataset of proteases. The clustering based on the dynamics adds interesting information to that known on the basis of structural and evolutionary relationships between the members of the protein family, thus facilitating the interpretation of dynamics as a bridge between protein structure and function. In addition, we used NMA and the *β*-GNM to build a basis set of vectors of the highdimensional space of the PA clan large-scale dynamics, and tested the basis set to demonstrate that it is sufficiently complete to describe the main large-scale dynamical features of the members of the dataset. The basis set of conformational motions was also successfully validated by comparison with results from MD simulations of proteins internal and external to the initial dataset.

In this regard, the method proved to deal particularly well with the conformational dynamics of structured regions; loops and disordered regions are by definition challenging to describe with an ENM, which is able to reproduce only small-amplitude fluctuations with respect to a well-defined reference structure; the dynamics of such regions, however, is qualitatively different from the functional one of the structured part, which is the one responsible to carry out the biological function in the proteins under examination. Additionally, we note that the dataset employed contained only a number of proteins belonging to the family of chymotrypsin-related proteases: a larger dataset is expected to lead to more general results; however, the number of proteins included was limited by the availability of experimental structures and by the choice to remove proteins with too high sequence identity. The natural development of the methodology presented and discussed in this work is its application to a larger dataset of proteins, comprehensive of multiple enzyme superfamilies, with the aim of building a basis set of conformational motions that represents a general vocabulary of proteins’ common dynamics. The method can then be employed to identify those common structural signatures that characterise the dynamics encoded in the basis components and relate them to specific biological functions.

## Author contributions

Conceptualization: R.P.; methodology, data collection and analysis: T.T. and M.R.; writing—original draft preparation: T.T. and G.M.; writing—review and editing: T.T., G.M., M.R. and R.P.; supervision: R.P.; funding acquisition: R.P. All authors have read and agreed to the published version of the manuscript.

## Data availability

The raw data produced and analysed in this work are freely available on the Zenodo repository at https://doi.org/10.5281/zenodo.6669245.

## Acknowledgments

The authors thank Roberto Menichetti for a critical and insightful reading of the manuscript. This project received funding from the European Research Council (ERC) under the European Unions Horizon 2020 research and innovation program (Grant 758588).

