## Supplemental material for "In search of a dynamical vocabulary: a pipeline to construct a basis of shared traits in large-scale motions of proteins"

### S1 Additional figures

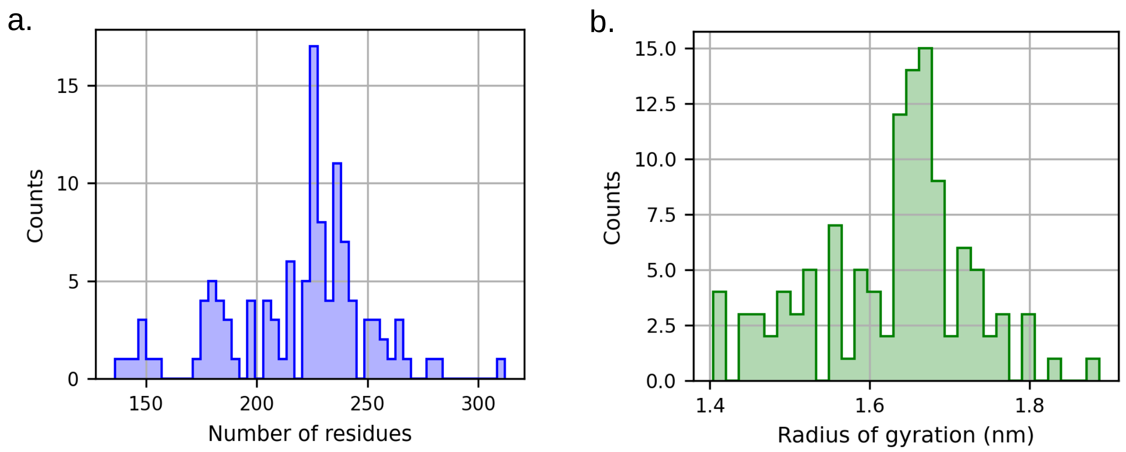


Figure S1: Histograms of the sequence length (**a**) and radius of gyration (**b**) of the proteins in the dataset.

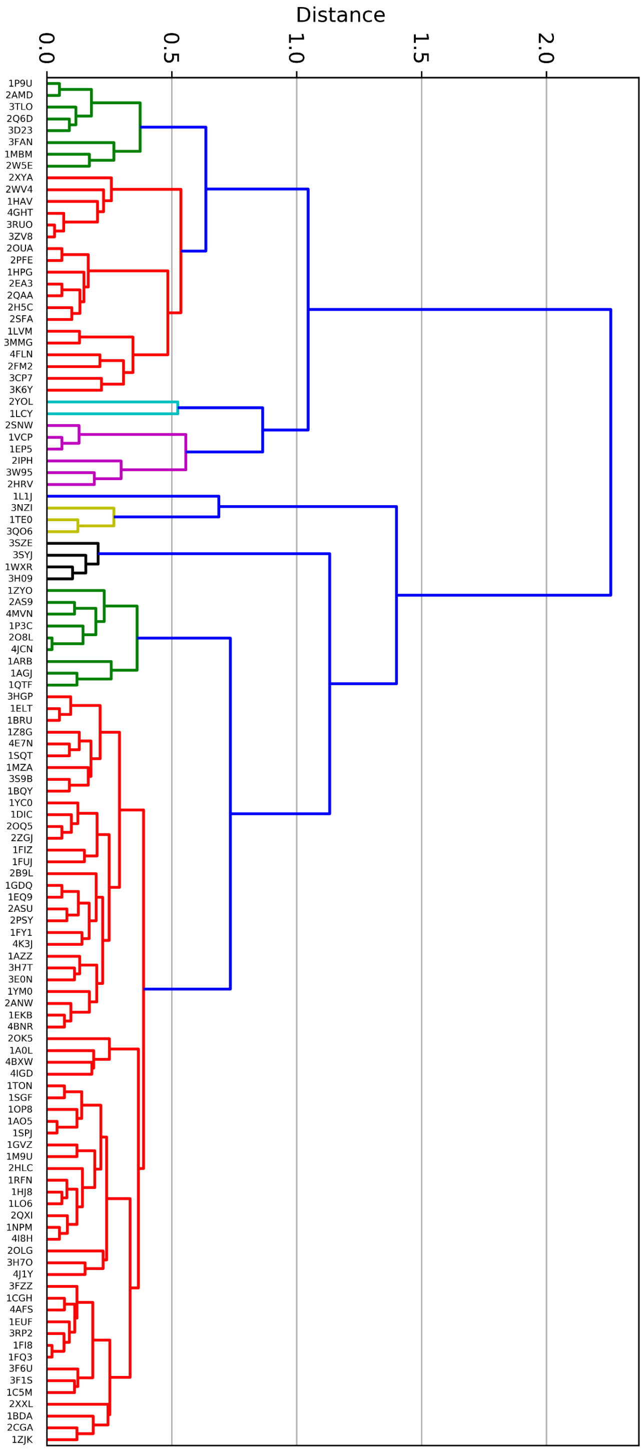


Figure S2: Dendrogram resulting from the hierarchical clustering, performed on the basis of the distance in dynamics between the dataset elements. The labels represents the PDB IDs, and colors are used to differentiate the clusters.

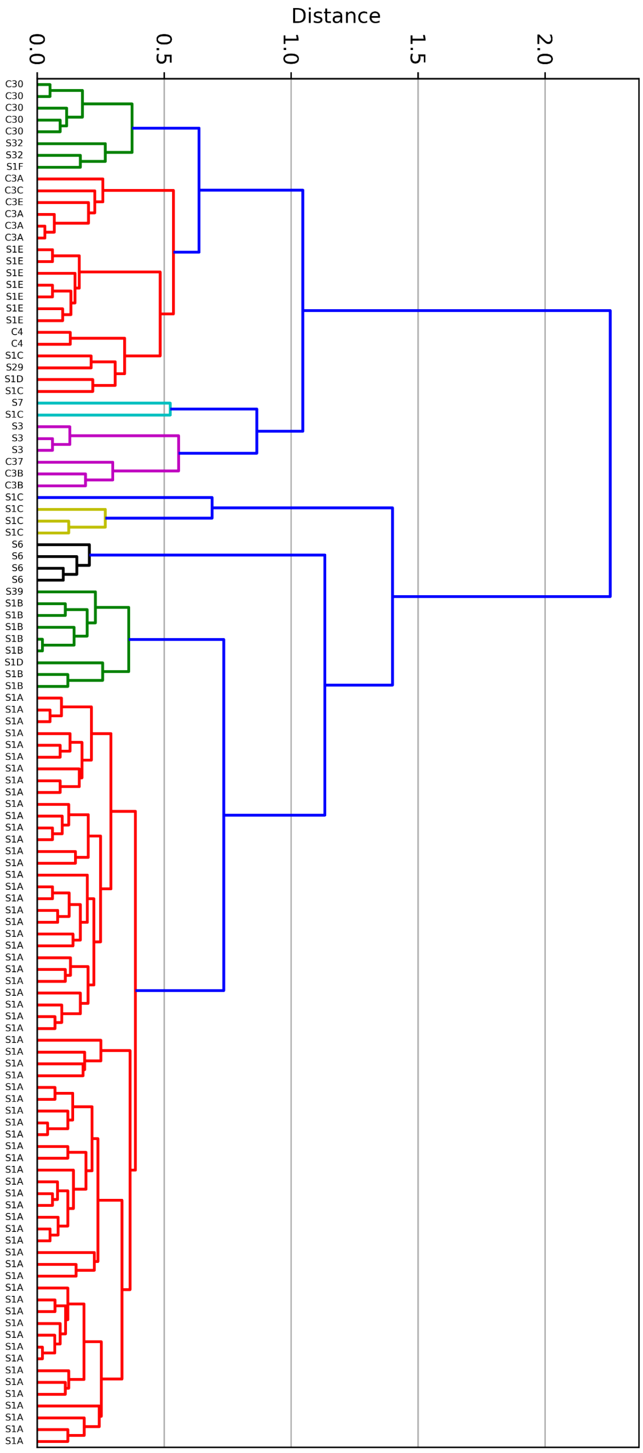


Figure S3: Dendrogram resulting from the hierarchical clustering, performed on the basis of the distance in dynamics between the dataset elements. The labels represents the protease subfamily of each protein, and colors are used to differentiate the clusters.

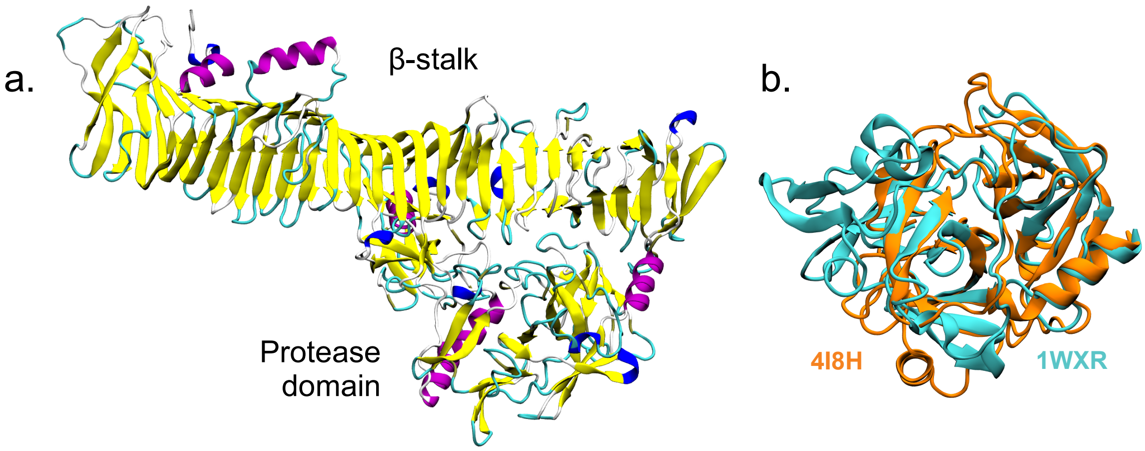


Figure S4: **a.** Full structure of the 1WXR protease from subfamily S6, displaying the long $\beta$-stalk domain at the C-terminus. **b.** Structural alignment of 1WXR (in cyan) and 4I8H from subfamily S1A (in orange), showing the similarity of their protein core.

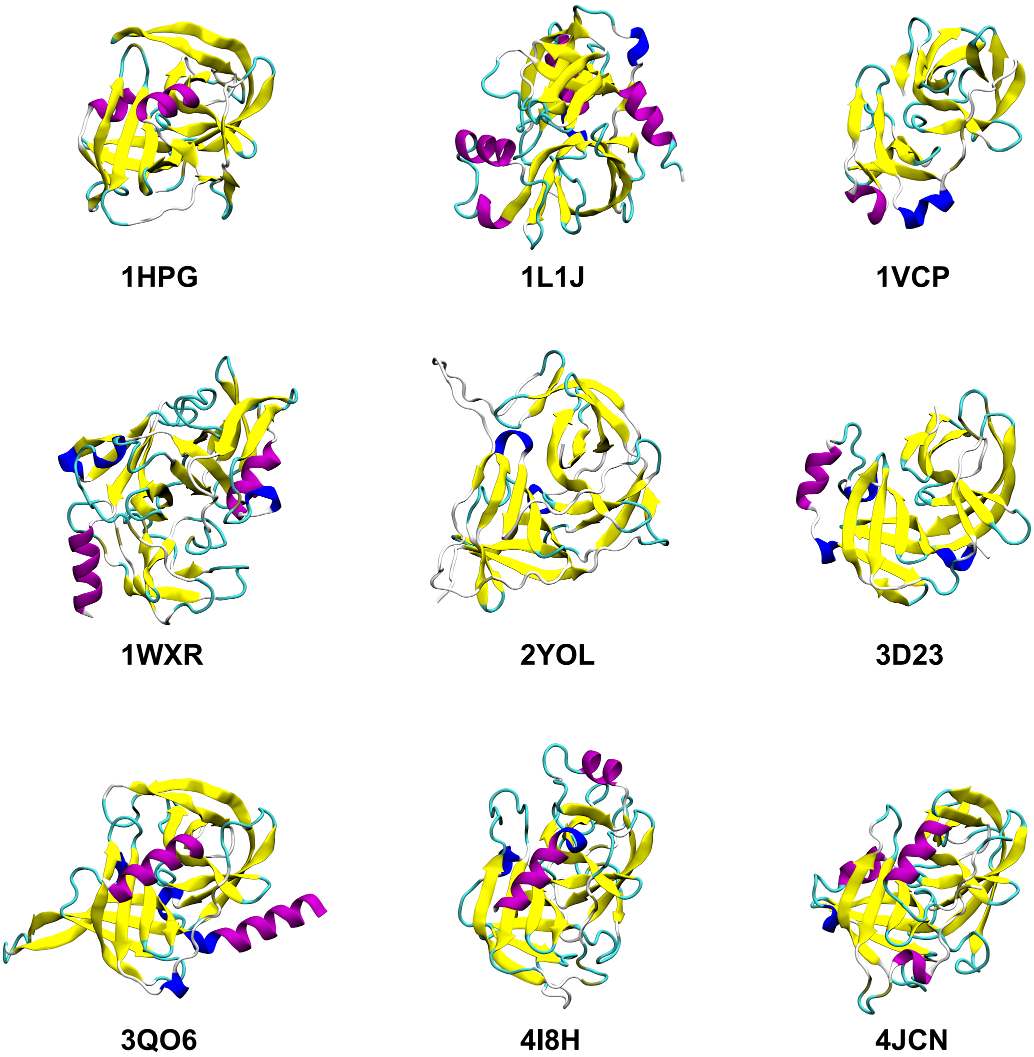


Figure S5: Structure of the representatives of each protein cluster, colored according to the secondary structure.

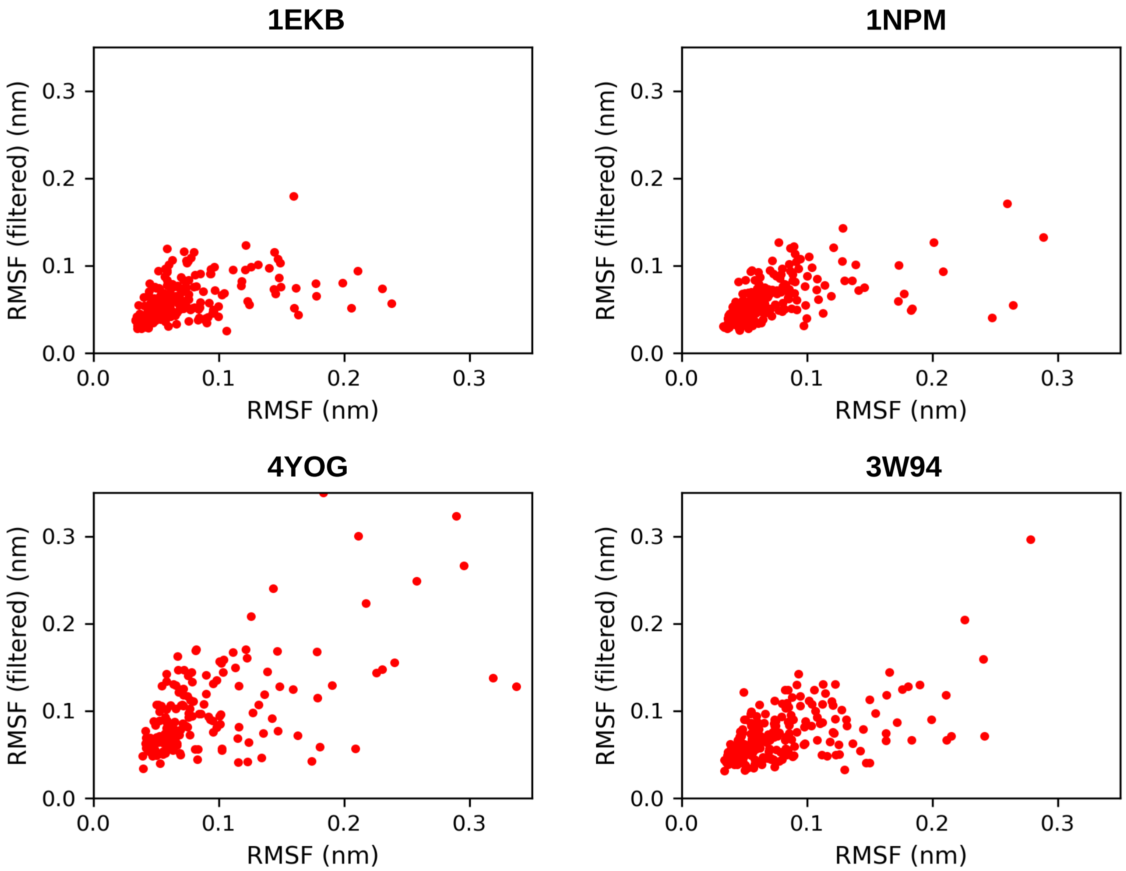


Figure S6: Scatter plots of the root-mean-square fluctuation values, computed on the $C_\alpha$ atoms.

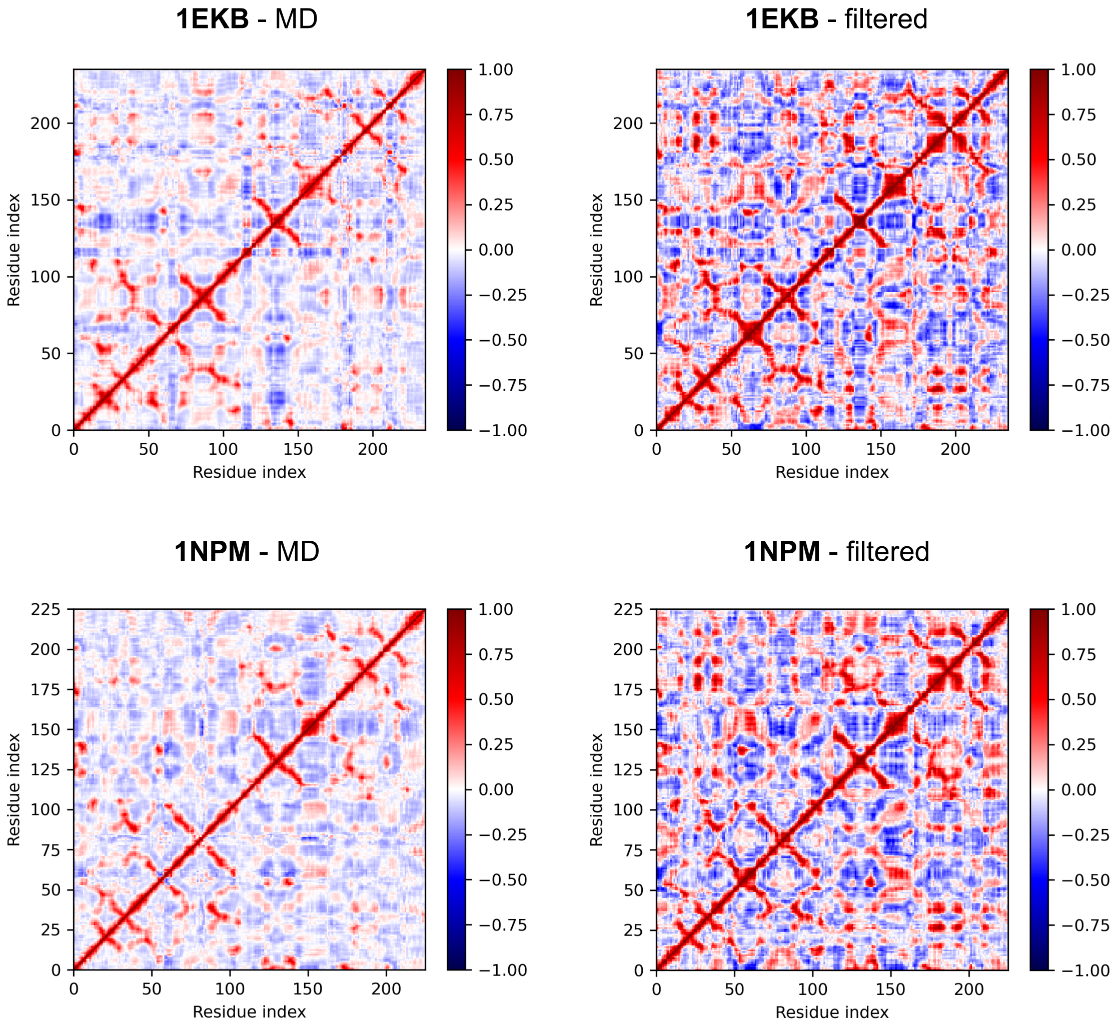


Figure S7: Cross-correlation computed from the simulations of the proteins 1EKB and 1NPM, both on the original and filtered trajectories. Both proteins belong to the dataset.

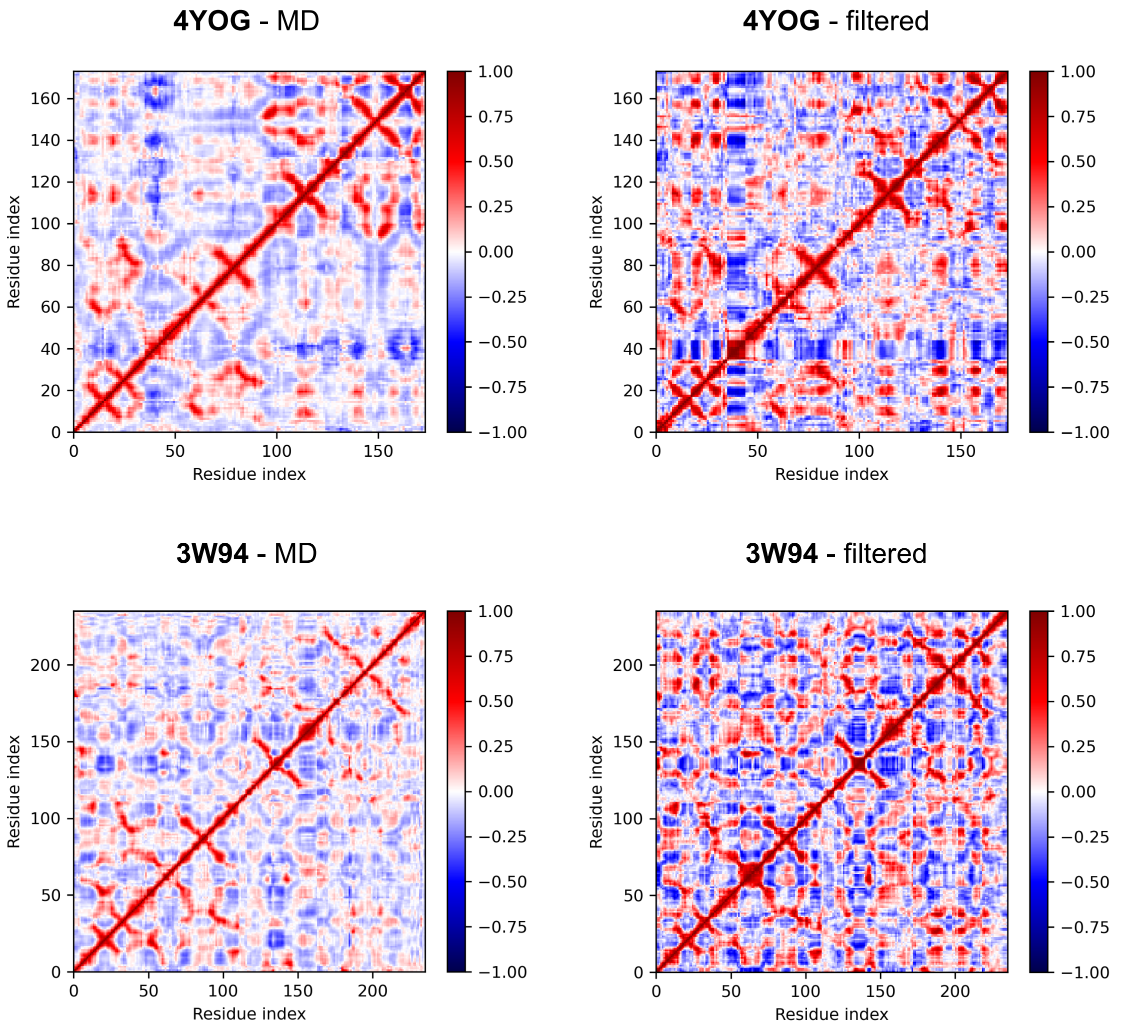


Figure S8: Cross-correlation computed from the simulations of the two proteins 4YOG and 3W94, both on the original and filtered trajectories. The two proteins are not part of the dataset from which the basis set is derived.

### S2 Additional tables

Table S1: List of the PDB IDs of the proteins comprising the dataset.

| |
|---|
| 1A0L 1CGH 1FY1 1MBM 1TE0 2AS9 2O8L 2SNW 3F1S 3QO6 |
| 4BXW 1DIC 1GDQ 1MZA 1TON 2ASU 2OK5 2W5E 3F6U 3RP2 |
| 4E7N 1EKB 1GVZ 1NPM 1VCP 2B9L 2OLG 2WV4 3FAN 3RUO |
| 4FLN 1AGJ 1ELT 1HAV 1OP8 1WXR 2CGA 2OQ5 2XXL 3FZZ |
| 3S9B 4GHT 1AO5 1EP5 1HJ8 1P3C 1YC0 2EA3 2OUA 2XYA |
| 3H09 3SYJ 4I8H 1ARB 1EQ9 1HPG 1P9U 1YM0 2FM2 2PFE |
| 2YOL 3H7O 3SZE 4IGD 1AZZ 1EUF 1L1J 1QTF 1Z8G 2H5C |
| 2PSY 2ZCH 3H7T 3TLO 4J1Y 1BDA 1FI8 1LCY 1RFN 1ZJK |
| 2HLC 2Q6D 2ZGJ 3HGP 3W95 4JCN 1BQY 1FIZ 1LO6 1SGF |
| 1ZYO 2HRV 2QAA 3CP7 3K6Y 3ZV8 4K3J 1BRU 1FQ3 1LVM |
| 1SPJ 2AMD 2I6Q 2QXI 3D23 3MMG 4AFS 4MVN 1C5M 1FUJ |
| 1M9U 1SQT 2ANW 2IPH 2SFA 3E0N 3NZI 4BNR |
